# Using asymmetry to your advantage: learning to acquire and accept external assistance during prolonged split-belt walking

**DOI:** 10.1101/2020.04.04.025619

**Authors:** Natalia Sánchez, Surabhi N. Simha, J. Maxwell Donelan, James M. Finley

## Abstract

People often adapt their coordination patterns during walking to reduce energy cost by using sources of external assistance in the environment. Adaptation to walking on a split-belt treadmill, where one belt moves faster than the other, provides an opportunity for people to acquire positive work from the treadmill to reduce metabolic cost by modifying where they step on the faster belt. Though we know what people should do to acquire this assistance, this strategy is not observed during typical adaptation studies. Here, by extending the duration of adaptation, we show that people continuously optimize energetic cost by adjusting foot placement to acquire positive work from the treadmill and reduce the work performed by the legs. These results demonstrate that learning to acquire and take advantage of assistance to reduce energetic cost is central in shaping adaptive locomotion, but this process occurs over timescales longer than those used in typical studies.

## Introduction

Humans frequently adapt their locomotor patterns to take advantage of potential sources of external assistance. For example, we use the pull on the leash from our dog to propel us as we walk down the street, and we use gravity when walking downhill to reduce the mechanical work performed by our muscles and reduce metabolic cost (Hunter et al., 2010). In other settings, we actively adjust our position in the environment to gain external assistance. During surfing, we kick and paddle to position our bodies and the surfboard to allow the waves to propel us. During sailing, we adjust the alignment of the sails relative to the wind to propel us through the water. Based on this tendency for the neuromotor system to take advantage of external assistance, several devices have been used to provide a source of external assistance and give the user the opportunity to reduce the work performed by the legs and the energetic cost of walking (Collins et al., 2015; Ding et al., 2018; Kang et al., 2019; Kim et al., 2019; Malcolm et al., 2013; Sawicki et al., 2020; Sawicki and Ferris, 2008; Zhang et al., 2017).

The success of assistive devices for reducing energetic cost depends greatly on the neuromotor system’s ability to adapt coordination to take advantage of assistance. First, the person must sense the points during the gait cycle when the device is providing assistance. Then, in some cases, the person must adjust their coordination patterns to acquire the assistance (Sawicki and Ferris, 2008). Then the person must accept the assistance provided by the external system by adapting the amplitude and timing of muscle activity at specific points in the gait cycle, to reduce muscular work and metabolic cost. These adjustments must take place while continuing to achieve high-level objectives, such as maintaining a steady walking pace when walking on a treadmill. Understanding how people learn to take advantage of assistance can provide novel insights into the process by which people optimize energetic cost during locomotion.

We recently demonstrated that a common task used to study locomotor learning, split-belt treadmill adaptation (Dietz et al., 1994; Reisman et al., 2005), provides an opportunity where people can learn to take advantage of external assistance provided by the treadmill. Specifically, they can gain assistance in the form of net positive work from the treadmill if they redistribute braking and propulsive forces between the two belts such that the leg on the fast belt produces most of the braking, and the leg on the slow belt produces most of the propulsion (Sánchez et al., 2019). The person can then take advantage of the work performed by the treadmill to reduce the positive work performed by the person’s muscles and ultimately reduce metabolic cost. We empirically tested these predictions and showed that when people use guided feedback to increase braking by stepping further forward with the leg on the fast belt, the treadmill performs net positive work on the person. This strategy led to asymmetric step lengths, where fast and slow step lengths are defined as the fore-aft distance between the feet at the respective foot-strike. This change in asymmetry was accompanied by reductions in the positive work generated by the legs and an associated reduction in metabolic cost (Sánchez et al., 2019). These findings complement previous studies showing that people reduce the mechanical work generated by the fast leg (Selgrade et al., 2017a, 2017b) and the metabolic cost of walking (Buurke et al., 2018; Finley et al., 2013) as they adapt to the split-belt treadmill.

Although people can take advantage of the positive work performed by the treadmill by adopting asymmetric step lengths, the vast majority of studies of split-belt adaptation have reported that adaptation ends with people taking steps of nearly equal length (Day et al., 2018; Leech and Roemmich, 2018; Long et al., 2016; Malone et al., 2012; Mawase et al., 2017, 2012; Reisman et al., 2005; Roemmich et al., 2016; Torres-Oviedo and Bastian, 2010). We recently showed that after people are provided with guided experience of less energetically costly step length asymmetries, they adopt an asymmetric gait pattern when they are allowed to adapt freely. This pattern consists of longer step lengths with the leg on the fast belt (Sánchez et al., 2019). It remains to be determined why people do not typically walk with longer steps with the leg on the fast belt during typical adaptation experiments. One potential explanation is that conventional adaptation studies may simply be too short in duration. Consistent with this idea, people took longer steps with the leg on the fast belt after 45 minutes of guided experience walking on the treadmill (Sánchez et al., 2019), which is three to four times longer than the length of conventional adaptation studies. People also take longer steps with the leg on the fast belt after multiple sessions of adaptation over five days (Leech et al., 2018), totaling approximately one hour and fifteen minutes of experience. Thus, the adoption of a gait with longer steps with the leg on the fast belt, which is less energetically costly, seems to occur after combined exposure times that are several multiples of the traditional 10-15 minutes allotted for adaptation.

Energy optimization during motor learning is a complex problem, which often requires extended experience. Although we can reduce energy cost over short timescales commonly used in studies of adaptation (Buurke et al., 2018; Finley et al., 2013; Huang et al., 2012; Huang and Ahmed, 2014; Selinger et al., 2019, 2015), energetic cost can be further reduced over multiple sessions of experience (Lay et al., 2002; Sawicki and Ferris, 2008; Sparrow and Newell, 1994). Adjusting movement patterns to reduce energetic cost poses a complex problem for the neuromotor system. Specifically, during split-belt walking people can reduce energy by learning to acquire external assistance in the form of positive work from the treadmill and then accept this assistance by reducing the positive work by the legs. This process requires adjusting interlimb coordination to redistribute the amount of work generated by each leg as well as adjusting the time during the gait cycle when the legs generate work (Selgrade et al., 2017a). To remain in place on the treadmill, the net mechanical work on the person must be zero on average. Therefore, if the person adopts a gait pattern that leads to an increase in the amount of positive work by the treadmill on the body, this must be balanced in one of three ways. The person can either dissipate the energy transferred to the body by increasing the negative work by the legs, reduce the positive work generated by the legs, or use a combination of both strategies. The person can fulfill this requirement with one leg or with both legs and they could employ these strategies at different points of the gait cycle. The challenge of both acquiring and accepting assistance in a way that leads to a net energetic benefit might explain why previous studies have not observed longer steps with the leg on the fast belt—in these studies, learning may simply have been incomplete.

In this study, we have two goals. The first goal is to understand whether extending the duration allowed for adaptation leads people to adopt asymmetric step lengths during split-belt walking. The second goal is to study the time course of how people learn to acquire positive work from the treadmill and the time course of how they learn to accept that work by reducing the positive work done by the legs. We hypothesized that extending the duration of experience on a split-belt treadmill would result in participants adopting asymmetric step lengths to increase the assistance they can acquire from the treadmill. During the extended exposure time, participants would also learn to accept this assistance and reduce the positive work generated by the legs. We also hypothesized that continued reductions in positive work by the legs would lead to continued reductions in metabolic cost beyond the cost of walking with steps of equal length. If our findings are consistent with these predictions, this will support the idea that walking with symmetric step lengths is only a point in the process by which people learn to take advantage of the assistance provided by the treadmill during split-belt adaptation. Importantly, this would suggest that locomotor adaptation requires significantly longer than 10-15 minutes of experience, likely because of the challenge associated with learning to take advantage of assistance. Finally, if participants can adjust their coordination to take advantage of external assistance and reduce the work generated by the legs, this would provide additional support for the role of energy optimization in shaping adaptation of locomotor behaviors.

## Materials and Methods

### Experiment Design

A convenience sample of fifteen healthy young adults participated in our study. Exclusion criteria included a history of lower extremity surgery or orthopedic injury within the last two years. All participants were naïve to the split-belt protocol, and right leg dominant, as assessed by self-reports. One participant terminated the experiment after the split-belt adaptation trial. The University of Southern California Institutional Review Board approved all experimental procedures, and each participant provided written informed consent before testing began. All aspects of the study conformed to the principles described in the Declaration of Helsinki.

We calibrated the motion capture system immediately before each experiment per manufacturer specifications (Qualisys AB, Goteborg, Sweden), including zeroing of the force plates embedded in the dual belt treadmill (Fully Instrumented Treadmill, Bertec Corporation, OH). We allowed the metabolic cart used to measure metabolic cost to warm up for thirty minutes and then we calibrated it per manufacturer specifications (Parvomedics, UT).

Participants completed three different walking trials. Participants first walked on the treadmill with both belts moving at 1.0 m/s for six minutes. We measured baseline spatiotemporal variables and metabolic cost during this trial (Fig. 1A – C). Participants then completed a continuous 45-minute split-belt adaptation trial. We collected data continuously for the entirety of the trial. Participants did not take any breaks during the adaptation trial. We set the speed of the left belt at 1.5 m/s, and the speed of the right leg at 0.5 m/s in a 3:1 ratio (Fig 1C). Participants then completed a 10-minute washout trial with belts tied at 1.0 m/s.

**Figure 1.**
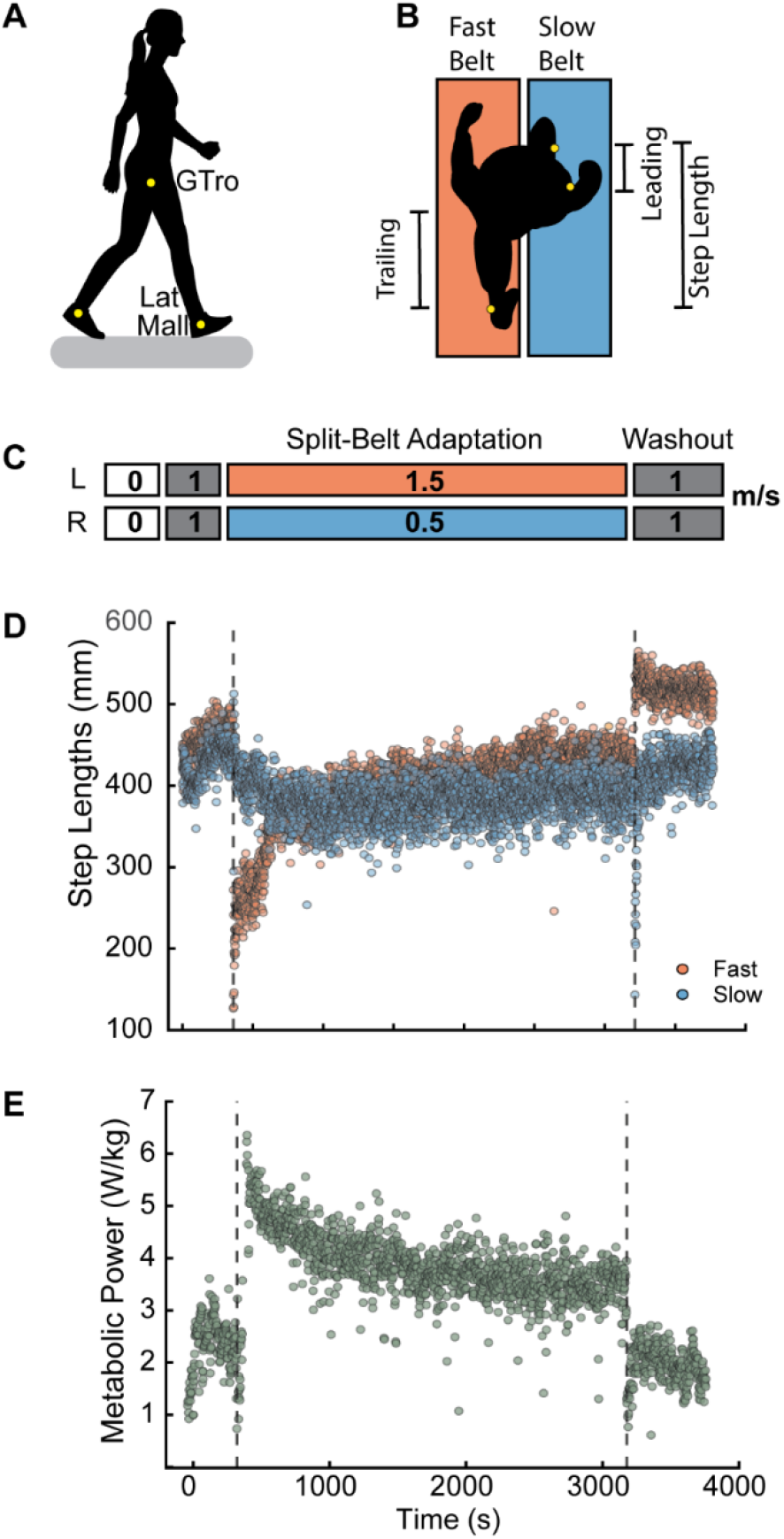
Experimental Design. A) Experimental setup. Participants walked on an instrumented, dual belt treadmill. B). Top view of the experimental setup and the leading and trailing limb placement relative to the body that make up individual step lengths. C) Experimental protocol and treadmill speeds for each trial. Participants rested for at least five minutes between all walking trials, to ensure metabolic cost was at standing baseline conditions before starting a new trial. D) Raw values of step lengths calculated using the distance between markers on the lateral malleoli for a single participant during all walking trials. Vertical dashed lines indicate different walking trials. Red: step lengths at heel-strike on the fast belt. Blue: step lengths at heel-strike on the slow belt. E) Raw values of net metabolic power for a single participant. Vertical dashed lines indicate different walking trials.

During all walking trials, participants wore a harness designed to prevent falls while providing no body weight support. No handrails were accessible during the experiment and participants were instructed to avoid holding on to the harness. After each walking trial, participants rested for at least four minutes, and we used real-time measures of metabolic cost to ensure that participants’ metabolic cost returned to resting levels before beginning the next walking trial.

### Data Acquisition

We recorded the positions of reflective markers located bilaterally on the lateral malleoli and greater trochanters at 100 Hz (Fig. 1A). We also recorded ground reaction forces generated by each leg at 1000 Hz to calculate mechanical power and work. We assessed metabolic cost by determining the rates of oxygen consumption 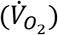 and carbon dioxide production 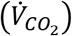 using a TrueOne^®^ 2400 metabolic cart (Parvomedics, UT) (Fig. 1E). The metabolic cart sampled gas concentrations on a breath-by-breath basis.

### Data Processing and Analysis

We used custom-written code in MATLAB R2019b (Mathworks, Natick, MA) for all data processing and analyses. A fourth-order low-pass digital Butterworth filter smoothed marker data and ground reaction force data. The filter cut-off frequency for marker data was set to 10 Hz and the filter cut-off frequency for force data was set to 20 Hz.

We used the positions of markers on the lateral malleoli (Fig. 1A) to estimate step length asymmetry as follows (Choi et al., 2009; Reisman et al., 2005; Roemmich et al., 2016):

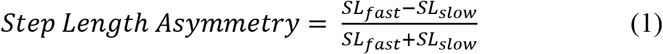

Here *SL*_*fast*_ is the step length at leading leg heel-strike on the fast belt, and *SL*_*slow*_ is the step length at leading leg heel-strike on the slow belt (Fig. 1B – D). Negative values correspond to longer steps with the slow (right) leg and positive values correspond to longer steps with the fast (left) leg. Step lengths are the product of adjusting the leading and trailing limb placement at heel-strike and toe-off respectively. Limb placement will affect the fore-aft component of the ground reaction force and the amount of work the legs must generate. Therefore, we calculated leading and trailing limb placement for each limb as the fore-aft distance relative to the midpoint between markers placed bilaterally on the greater trochanters (Fig 1B), to determine whether individuals adjusted limb placement in a manner consistent with taking advantage of the work by the treadmill.

We also calculated stance, swing and double support times from marker data (Zeni et al., 2008). Swing time corresponds to the time between toe-off, which was estimated as the most posterior location of the ankle markers, to heel strike on the same side, which was estimated as the most anterior location of the ankle marker. Stance time corresponds to the time between heel strike and toe off on the same side. Finally, double support time for a given limb corresponds to the time from contralateral heel strike to ipsilateral toe off.

We estimated the mechanical work performed by the legs using an extension of the individual limbs method (Donelan et al., 2002; Sánchez et al., 2019; Selgrade et al., 2017b, 2017a). This method considers a person as a point mass body with legs that are massless pistons. The legs can generate forces on the ground while simultaneously generating equal but opposite forces on the point mass, referred to as the center of mass. Thus, we assume that all the work performed by the person is being performed by the legs. To calculate mechanical work in our experiments, we segmented force data into strides using a vertical ground reaction force threshold of 32N to identify foot strike (Selgrade et al., 2017a) and then calculated the center of mass acceleration in the fore-aft, vertical, and medio-lateral directions, as the sum of the forces within a stride normalized by body mass. We then calculated the center of mass velocities in each direction as the time integral of the accelerations. We determined the integration constant for each stride by forcing the average velocity over the stride to be zero in each direction. All velocities and accelerations were expressed relative to a reference frame attached to the fixed ground. Next, we calculated the instantaneous power generated by each leg for each stride as the instantaneous sum of the dot product of the ground reaction force and the center of mass velocity and the dot product of the force applied to the respective belt and the belt speed. Finally, we calculated the total positive and total negative work performed by each leg as the time integral of the positive or negative portion of the total instantaneous power over the stride cycle. To have a comparable interpretation with metabolic power, we express all measures of work as work rate by dividing each measure by stride duration.

We estimated the energy consumed using the standard Brockway equation (Brockway, 1987) as follows:

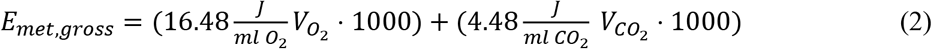

From here, we divided *E*_*met,gross*_ by the exact duration (T) over which it was calculated to obtain an estimate of the gross metabolic rate *P*_*met,gross*_ measured in Watts. Finally, we subtracted each participants’ standing metabolic rate from each walking trial (Fig 1E). Thus, all metabolic rate values presented here are net metabolic rate. For each participant, we estimated metabolic cost as the average net metabolic rate of the three minutes preceding the timepoint of interest. This corresponded to the last three minutes of the standing baseline and baseline walking trials, and minutes 12 – 15 and 42 – 45 of adaptation.

Changes in step length asymmetry during conventional studies of split-belt adaptation occur over two distinct timescales (Darmohray et al., 2019; Mawase et al., 2012; Roemmich et al., 2016). However, it is unclear if two distinct timescales are necessary to model adaptation when the adaptation period is extended to 45 minutes. Therefore, we modeled step length asymmetry (SLA) as a function of stride number (*s*) for the adaptation and washout trials using either a single exponential (Equation 3) or the sum of two exponentials (Equation 4).

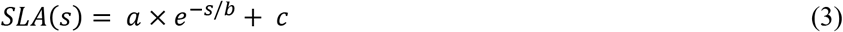

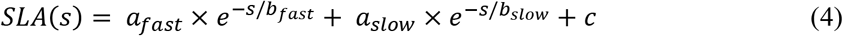

In the single exponential model, *a* corresponds to the step length asymmetry at the onset of the perturbation (*s* = 0), and *b* is the rate constant parameter. Based on our hypothesis, SLA would plateau at positive values, therefore, we added a term *c* which corresponds to the step length asymmetry as *s* → ∞. For the two-exponential model, *a*_*fas*t_ and *a*_*slow*_ are the initial values of the fast and slow exponentials respectively, and *b*_*fast*_ and *b*_*slow*_, are the fast and slow rate constants, respectively. We fit one and two exponential models to step length asymmetry data for the group as a whole by concatenating step length asymmetry vectors for all participants. We then used the Akaike information criterion (Akaike, 1981) to compare whether the more complex two-exponential model provided a better fit to the data than a single exponential model. We selected the model with the lowest AIC. Step length asymmetry data and the corresponding two exponential fits for individual participants are included in Supplementary Figure 1.

We used the exponential model selected above and ran bootstrap analyses to obtain the 95% confidence intervals of the adaptation model parameters. We created 10,000 bootstrapped samples of 15 participants by sampling participants with replacement. For each bootstrapped sample, we concatenated the *SLA* vectors for all participants and fit the data to the selected model using fitnlm in MATLAB. This process was repeated 10,000 times. We sorted all model parameters in ascending order and identified the 500^th^ and 9,500^th^ values as the limits of the 95% CI.

We also determined how the duration of adaptation influenced the magnitude and washout of after-effects associated with adaptation. We compared step length asymmetry for the first five strides of washout, which corresponds to early post-adaptation (Reisman et al., 2005). We analyzed washout data from our study in 14 participants and a previous study of 12 participants where they adapted for 15 minutes (Park and Finley, 2019, 2017). We implemented the models in equations 3 and 4 and the procedure described above to determine whether a one or two exponential model provided a better fit to the washout data. We also ran bootstrap analysis for the data from the washout period.

### Statistical Analyses

Our goal was to test the hypothesis that prolonged exposure to split-belt walking leads individuals to take longer steps with the leg on the fast belt, resulting in more positive step length asymmetries, increased positive work performed by the treadmill, reductions in positive work rate by the legs, and reductions in metabolic cost beyond those observed in traditional 15-minute adaptation experiments. Based on the Kolmogorov-Smirnov test all data were normally distributed. Therefore, we used paired samples, one-tailed t-tests for step lengths, step length asymmetry, foot placement relative to the body, stance, swing and double support times and the positive and negative mechanical work performed by the legs to test for differences between 15 and at 45 minutes of adaptation. For each variable, we averaged the values over the last 100 strides and the 100 strides before the 15-minute time-point to obtain values for statistical analyses.

We hypothesized that individuals would adopt positive step length asymmetries to gain assistance from the treadmill, which would allow them to reduce the positive work by the legs and reduce metabolic cost. Therefore, we computed the correlation between changes in step length asymmetry (Δ*SLA*_45−0_) and changes in positive work rate by the fast leg 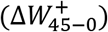 during adaptation to determine whether people decreased positive work by the fast leg as step length asymmetry becomes more positive. Similarly, we computed the correlation between the change in work 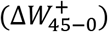 and the change in metabolic cost (Δ*MetCost*_45−0_) during adaptation because we hypothesized that people would reduce energetic cost as they reduced positive work. Given the rapid reductions in step length asymmetry that occur at initial exposure to the split-belt treadmill, for correlation analyses we calculated average step length asymmetry and work only for the first and last five strides (Reisman et al., 2005). For metabolic cost, we used data averaged over one minute for the early and late adaptation periods. The metabolic cost for early adaptation was averaged over minutes 3 – 4 of split-belt walking to account for transport lag and temporal dynamics of the change in metabolic cost in response to changes in exercise intensity (Finley et al., 2013; Selinger and Donelan, 2014). We used the Pearson correlation coefficient as data were normally distributed.

We compared step length asymmetry during early post-adaptation after 45 and 15 minutes of adaptation using independent samples t-tests. We also computed the effect size of the difference between model parameters for washout data after 45 minutes and after 15 minutes of adaptation using Cohen’s d (Cohen, 1992).

## Results

### After prolonged adaptation, people overshoot symmetry and adopt positive step length asymmetries

We hypothesized that extending the duration of experience on a split-belt treadmill would allow participants to step further forward with the leg on the fast belt, which would lead to positive asymmetries and allow the treadmill to generate net positive work on the person. In agreement with our hypothesis, participants continued to lengthen the step length on the fast belt, with an average increase of 18 mm from 15 to 45 minutes of adaptation (Fig 2A – C paired t-test one tail, 95% CI > 7.942, p=0.003), whereas the step length on the slow belt did not change systematically from 15 to 45 minutes of adaptation (Fig 2D – E, p=0.481). After 45 minutes of adaptation, step length asymmetry was on average 0.018 more positive compared to 15 minutes (Fig 2F – H, paired t-test, one tail 95% CI > 0.001, p=0.042). At 45 min, step length asymmetry was significantly more positive than baseline, with an average of 0.030 (Fig 2F, one sample t-test 95% CI> 0.004, p=0.020).

**Figure 2.**
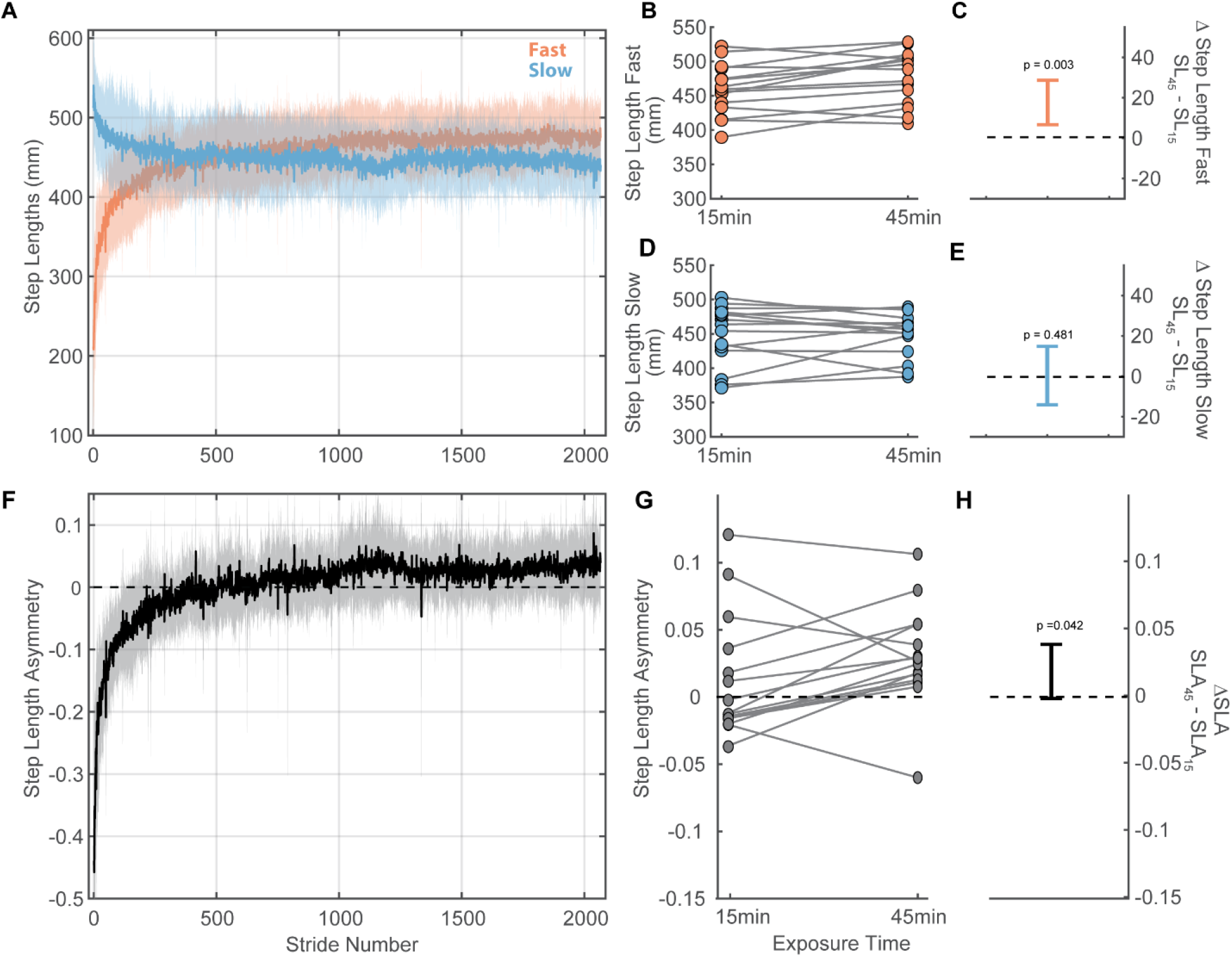
Participants continuously modified step lengths with the fast leg over 45 minutes of adaptation. A) Fast (red) and slow (blue) step lengths during adaptation. Solid lines represent sample means. Shaded areas are standard deviations. After 45 minutes of adaptation, the step length on the fast belt was longer than that on the slow belt (p=0.003). B) Average step lengths on the fast belt at 15 and 45 minutes for all participants. C) 95%CI of the differences in fast step lengths at 45 minutes compared to 15 minutes. D) Average step lengths on the slow belt at 15 and 45 minutes for all participants. E) 95%CI of the differences in slow step lengths at 45 minutes compared to 15 minutes. F) Mean step length asymmetry during adaptation (black). Grey shaded areas are standard deviations. G) Average step length asymmetry at 15 and 45 minutes for all participants. H) 95%CI of the differences in step length asymmetry at 45 minutes compared to 15 minutes. Averages in panels B, D and G were obtained for 100 strides. SLA: step length asymmetry. SL: step lengths.

Participants could take longer steps with the leg on the fast belt by either allowing the leg on the slow belt to trail further back or by placing the leg on the fast belt further forward at heel-strike. Based on our hypothesis, placing the leg further forward on the fast belt would increase the fore-aft component of the ground reaction force, allowing the treadmill to perform more positive work on the body. In agreement with this hypothesis, participants continued to adjust fast leg leading placement from 15 to 45 minutes by 13 mm on average (Fig 3A, C, paired t-test, one tail 95% CI > 6.464, p=0.003). Neither slow leg leading or trailing placement relative to the body (Fig 3B, D, F), nor fast leg trailing placement relative to the body changed systematically (Fig 3E). We excluded one participant from this analysis due to loss of greater trochanter marker data.

**Figure 3.**
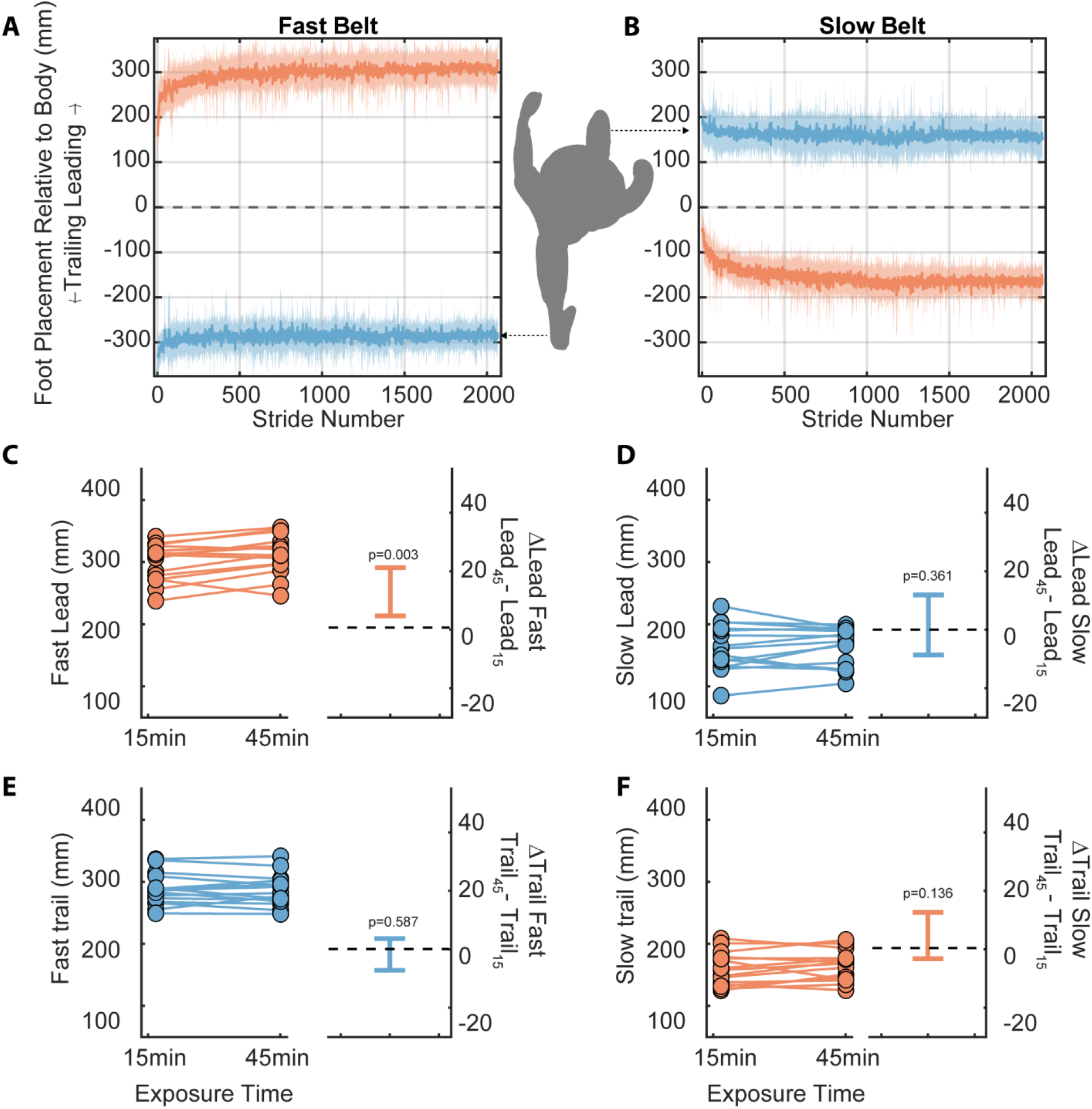
Modifications in foot placement at heel-strike and toe-off to increase fast step lengths are consistent with a strategy that would increase assistance gained from the treadmill. We present data for 14 participants as one participant lost a marker by the end of the trial. A) Fast belt. Top, fast leg placement at heel-strike. Bottom: fast leg placement at toe off. The increase in step length by the fast leg was due to further placement of leading limb at heel-strike. B) Slow belt. Top, slow leg placement at heel-strike. Bottom: slow leg placement at toe off. The distance between off-red curves indicates the fast step length. The distance between off-blue curves indicates the slow step length. Top view of participant included for visualization purposes. Shaded areas are standard deviations. C-F) Limb placement at 15 and 45 minutes and differences between the placement of the fast and slow limbs at 15 and 45 minutes.

Adaptation of step lengths occurred with a simultaneous adaptation of stance time and double support time. However, each of these temporal variables adapted over different timescales. Participant’s double support time plateaued quickly with no changes in fast (p=0.500) or slow (p=0.915) double-support times between 15 and 45 minutes. In contrast, participants’ stance time on the slow leg increased from 15 to 45 minutes by 0.03 ± 0.05 s (paired t-test, one tail 95% CI > 0.001, p=0.044), while stance time on the fast leg remained unchanged from 15 to 45 minutes (p=0.080). Accordingly, swing time on the fast leg was 0.02 ± 0.04 s longer at 45 compared to 15 minutes (CI > 0.001, p=0.049), whereas swing time on the slow leg remained unchanged (p=0.155). This indicates that participants also continue to adjust step timing over the prolonged split-belt adaptation.

### Participants quickly accept positive work from the treadmill, and gradually reduce positive work by the legs

Consistent with our hypothesis that prolonged exposure to the split-belt treadmill would allow participants to learn how to use the assistance provided by the treadmill, participants performed net negative work rate with the legs by the end of adaptation with an average of −0.06 W/kg (one sample t-test, one tail 95% CI < −0.0438, p=3.720×10^−6^). Net negative work rate by the legs was accomplished as follows. Participants increased negative work rate with the fast leg within the first hundreds of strides (Fig. 4A), with no significant changes in negative work rate from 15 minutes of adaptation to 45 minutes of adaptation for the fast leg (Fig 4A, 5A, paired t-test, p=0.094). The negative work rate by the slow leg did not change systematically between 15 minutes and 45 minutes of adaptation (Fig 4B, 5B, t-test p=0.087).

**Figure 4.**
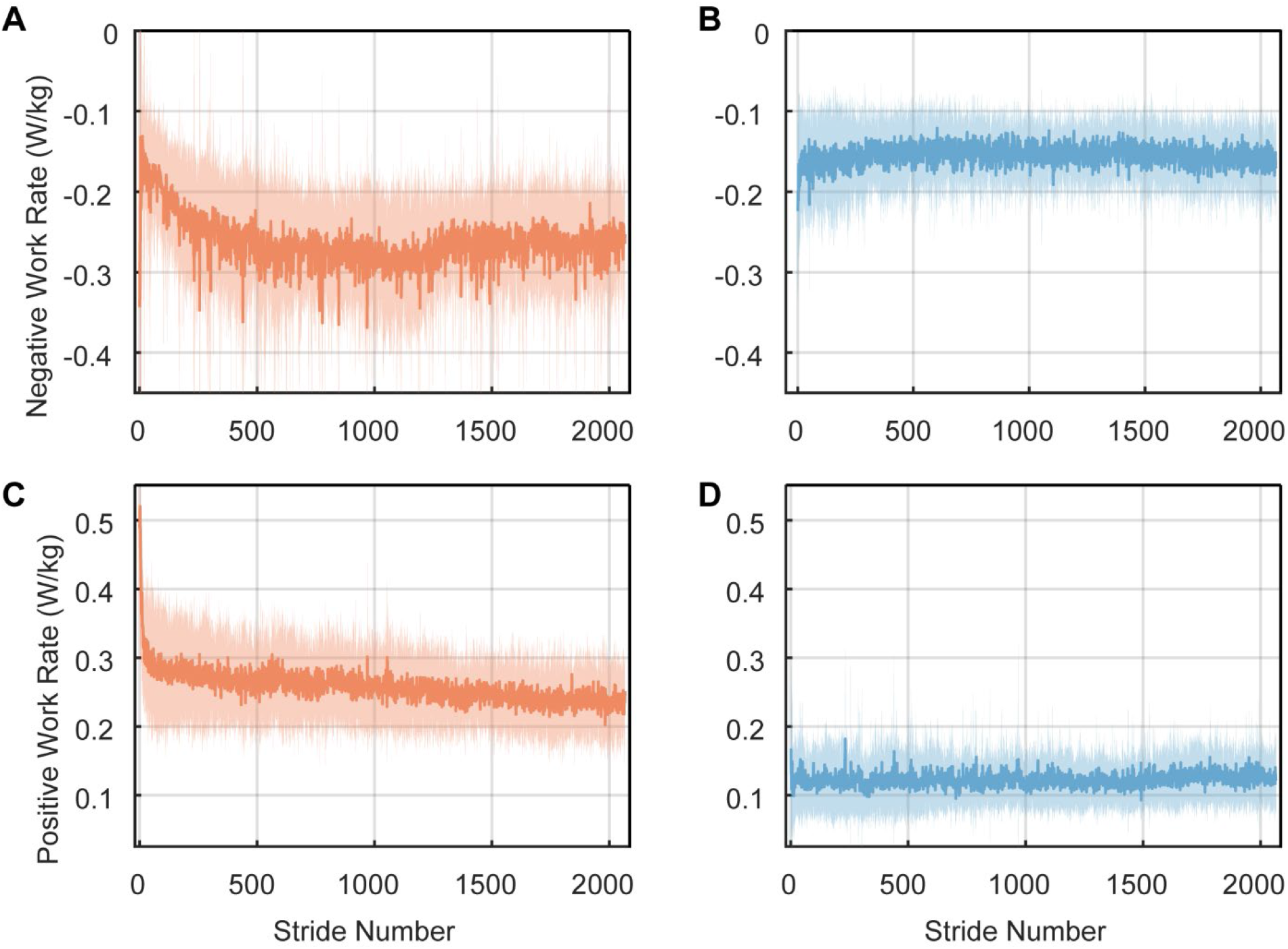
Mechanical work generated by the fast and slow leg continued to be adjusted during adaptation. A) Negative work rate generated by the fast leg. B) Negative work rate generated by the slow leg. C) Positive work rate by the fast leg. D) Positive work rate by the slow legs. Shaded areas are standard deviations.

**Figure 5.**
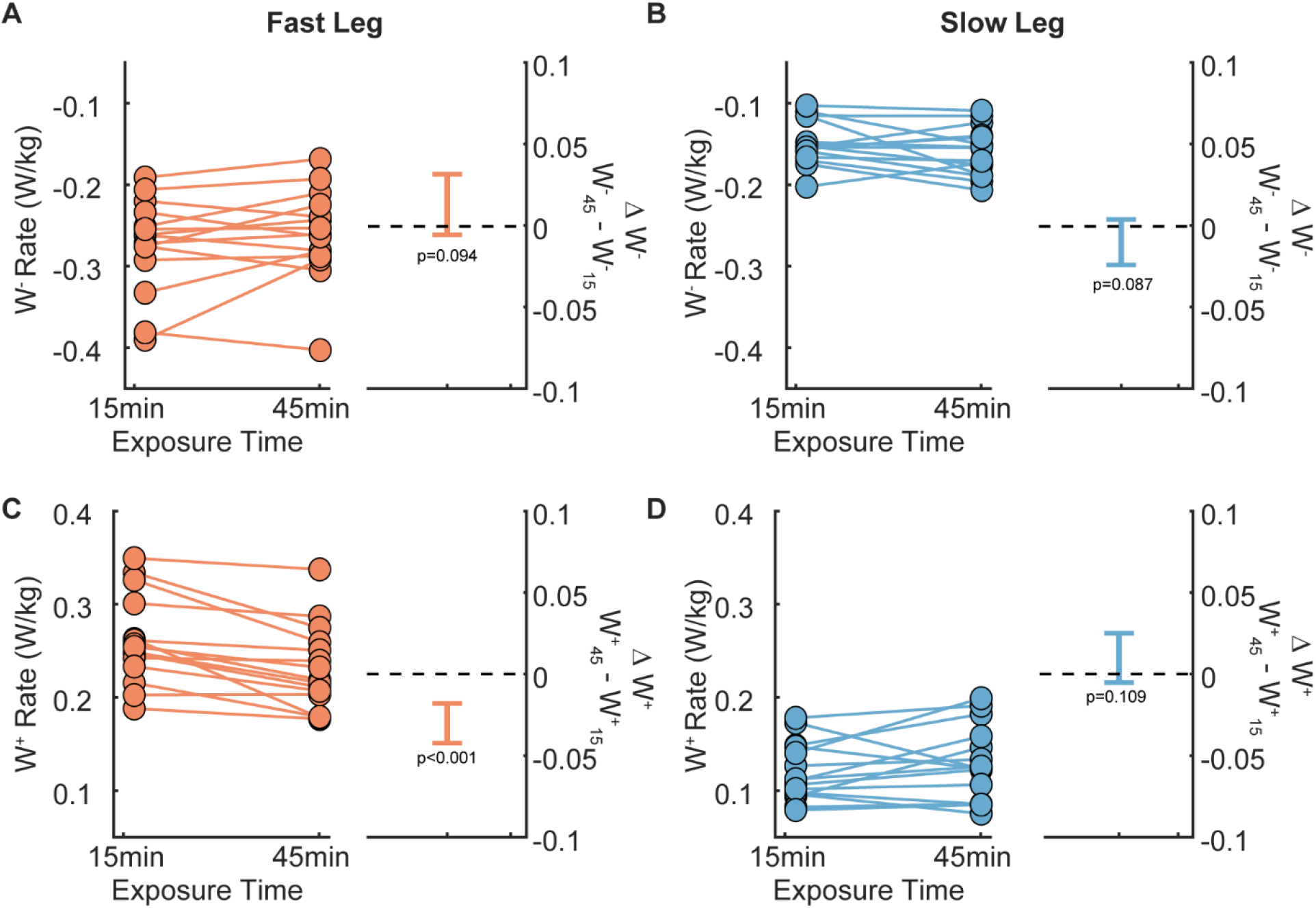
Changes in mechanical work rate by the fast and slow leg during prolonged exposure. A) Change in negative work rate for the fast leg. B) Change in negative work rate for the slow leg. No consistent changes in negative work rate were observed during prolonged exposure to the split-belt as seen in the 95%CI plots. C) Positive work rate by the fast leg continued to decrease throughout the 45 minutes of split-belt walking with an additional reduction from 15 to 45 minutes. D) Change in positive work rate by the slow leg. No changes in work rate by the slow leg were seen between 15 and 45 minutes.

To take advantage of the assistance provided by the treadmill, it is not enough to only increase negative work rate by the legs. Participants must also learn to decrease positive work rate, which could ultimately lead to a reduction in metabolic cost. Learning to reduce this positive work rate by the legs also begins quickly (Fig. 4C, Supplementary Fig. 2) with an average reduction from initial exposure to 15 minutes of adaptation of 0.20 W/kg (95% CI >0.16, p=1.14×10^−6^), or 42 ± 14% for the fast leg. This learning process continues throughout the entire experiment with an average reduction in positive work rate performed by the fast leg of 11 ± 9%, which corresponds to 0.025 W/kg from 15 to 45 minutes of (Figs. 4C, 5C, paired t-test, 95% CI> 0.019, p<1.48×10^−4^). Note that the reduction in positive work rate by the fast leg was not complete at the end of the experiment, as evidenced by the presence of a negative slope in the work rate timeseries. The positive work rate by the slow leg stabilized almost immediately with no changes between initial exposure and 15 minutes (Fig 4D, t-test, p=0.215) or between 15 and 45 minutes (Figs 4D, 5D, p=0.109).

We hypothesized that adoption of a more positive step length asymmetry would be associated with the reduction in positive work by the fast leg. In agreement with this hypothesis, we observed that changes in step length asymmetry during adaptation (Δ*SLA*_45−0_) were negatively correlated with changes in positive work rate 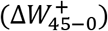 by the fast leg (r= −0.64, p=0.010).

### Metabolic cost continues to decrease as participants learn to take advantage of assistance from the treadmill

We also hypothesized that continuous reductions in the positive work rate by the legs over 45 minutes of adaptation would lead to reductions in the metabolic cost of walking beyond the cost of walking with steps of equal lengths. Consistent with this prediction, participants reduced their metabolic cost by an additional 7% (IQR 8%) which corresponds to a reduction of 0.17 W/kg from 15 to 45 minutes of walking (Fig. 6A – C, paired t-test 95% CI>0.059, p=0.007). The lower metabolic cost observed at 45 minutes indicates that people continue to refine their gait pattern to increase economy despite the demands of walking on a split-belt treadmill for 45 minutes.

**Figure 6.**
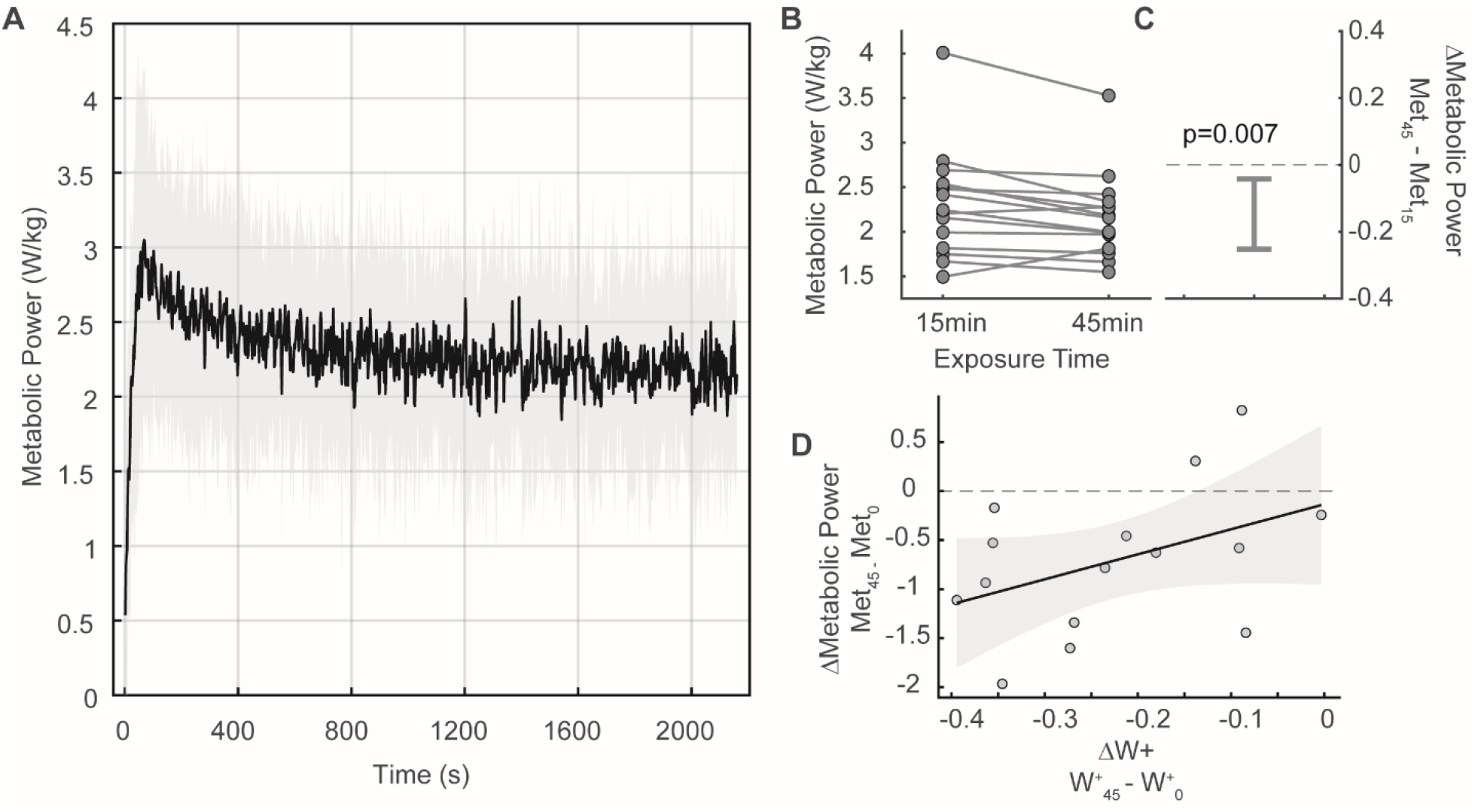
Metabolic power continuously decreased during adaptation. A) Average metabolic power during adaptation trial. Shaded area represents the standard deviation across participants. B) Average metabolic cost at 15 minutes and at 45 minutes of adaptation. Averages for metabolic data were obtained across three-minute periods. C) Reduction in metabolic cost from 15 to 45 minutes of adaptation. D) Correlation between the change in positive work by the fast leg from initial exposure to 45 minutes and change in metabolic power from initial exposure to 45 minutes.

In agreement with our hypothesis that walking with more positive step length asymmetry is less energetically costly, we found a negative correlation between the change in asymmetry (Δ*SLA*_45−0_) and the change in metabolic cost over the course of adaptation (Δ*MetCost*_45−0,_, r=−0.66, p=0.026). The change in metabolic cost measured during adaptation Δ*MetCost*_45−0_, was also positively correlated with the change in positive work rate by the fast leg, Δ*W*_45−0_ (r=0.52, p=0.048, Fig 6D). Thus, individuals reduced the positive work rate by the legs when obtaining assistance provided by the treadmill, and this ultimately led to reductions in metabolic cost.

### Adaptation of step length asymmetry can be characterized by a sum of two exponentials model that plateaus at positive step length asymmetries

A sum of two exponentials provided a good fit to the participant-level step length asymmetry data during adaptation (Supplementary Fig. 1). Results for the bootstrap analyses returned an R^2^ for the two exponential model of 0.356, bootstrap 95% CI [0.265, 0.447]. The model parameters were: *a*_*fast*_ = −0.31 (95% CI [−0.39, −0.24]), *a*_*slow*_ = −0.15 (95% CI [−0.20, −0.11]). *b*_*fast*_ = 20 strides (95% CI [9, 35]), and *b*_*slow*_ = 347 strides (95% CI [195, 553]). Finally, the step length asymmetry plateau, *c*, was equal to 0.033 (95% CI [0.015, 0.053]), which supports our hypothesis that participants adapt toward positive asymmetries. Based on this model, it would take approximately 1564 strides (95% CI [1089, 2060]) to adapt 95% of the difference between the initial step length asymmetry during early adaptation and the final plateau. This is on average twice as long as traditional adaptation experiments. Note that the average duration of our study was ~2300 strides (range between 2066 to 2853 strides for all 15 participants).

### The duration of washout is proportional to the duration of adaptation

We expected that when the belts returned to the same speed, individuals would have greater retention of the strategy adopted after 45 minutes of adaptation compared to 15 minutes, indicated by both a larger initial step length asymmetry and a longer time constant for washout. We did not observe differences in step length asymmetry during early post-adaptation after 45 minutes (0.34 ± 0.15) or 15 minutes (0.32 ± 0.08) of split-belt walking (Fig. 7A – B, independent samples t-test, p=0.758). However, the time course of washout varied with the duration of adaptation. A single exponential model best characterized the time course of changes in step length asymmetry during washout with average R^2^ values of 0.48 (95% CI [0.41, 0.56]) after 45 minutes of adaptation and 0.60 (95% CI [0.54, 0.66]) after 15 minutes of adaptation. The means and 95% confidence intervals of the model parameters were (Fig 7B – D): *a*_45_ = 0.22 (95% CI [0.18, 0.26]), *a*_15_ = 0.27 (95% CI [0.22, 0.31]), *b*_45_ = 63 strides (95% CI [41, 82]), *b*_*15*_ = 25 strides (95% CI [17, 33]), *c*_45_ = 0.05 (95% CI [0.035, 0.066]), and *c*_15_ = 0.038 (95% CI [0.034, 0.043]). Participants required almost three times longer to washout the learned pattern after 45 minutes versus 15 minutes of adaptation (Fig. 7A, C). The effect size for the difference in initial asymmetry (*a*_45_ *vs*. *a*_15_) was small with a Cohen’s d of 0.02. The difference in washout rates (*b*_45_ *vs*. *b*_15_) had a large effect size with a Cohen’s d of 13. Finally, the differences in the plateau (*c*_45_ *vs*. *c*_15_) had a small effect size with a Cohen’s d of 0.14.

**Figure 7.**
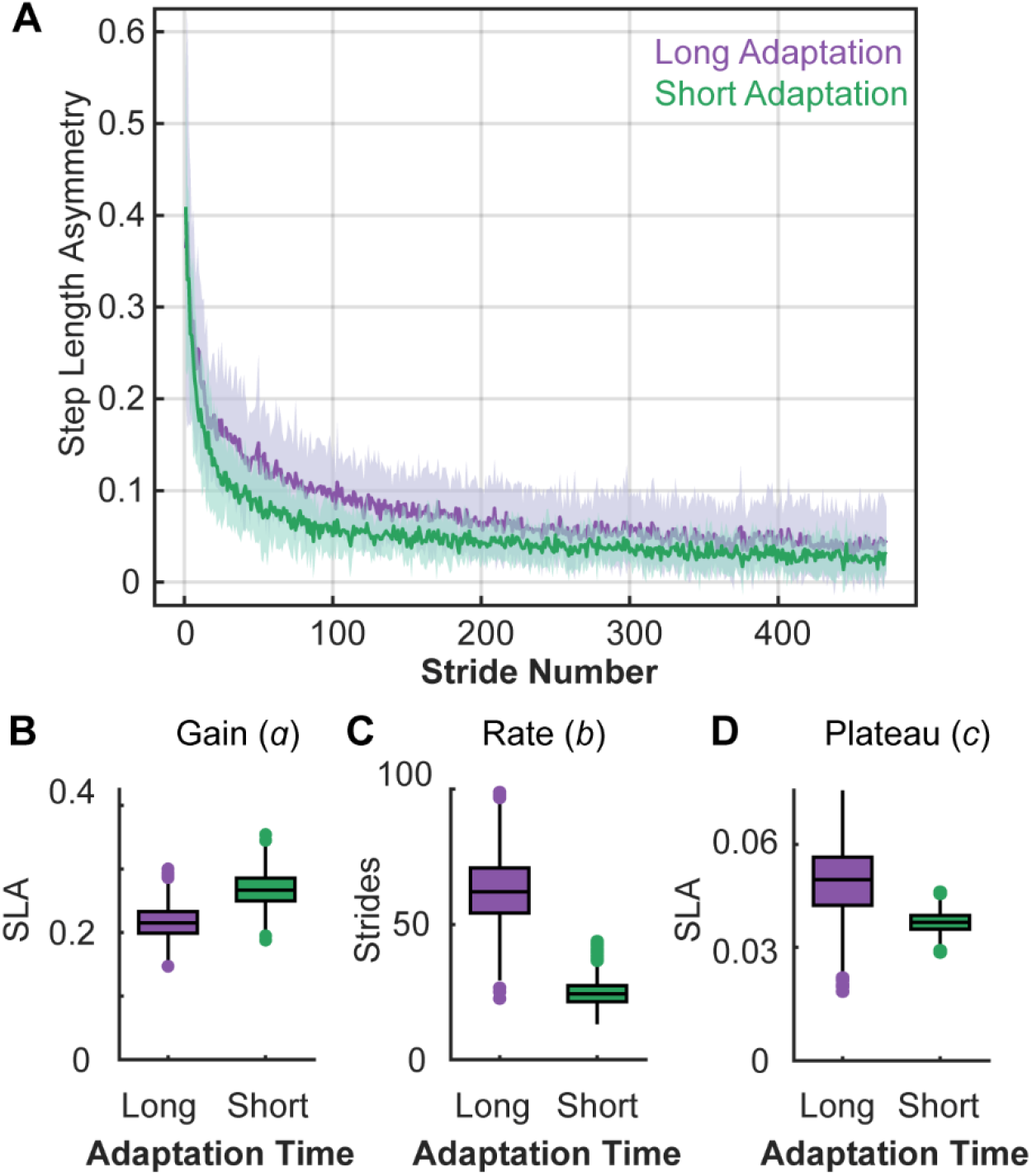
Step length asymmetry during washout trial. **A)** Step length asymmetry timeseries. Data in purple were collected in N=14 participants who completed our study. Data in green were collected from a previous study following a 15 minute adaptation period (Park and Finley, 2019). B-D) Distribution of the bootstrapped parameters for the single exponential model coefficients.

## Discussion

Learning to take advantage of external assistance from the environment during walking poses a complex problem for the neuromotor system. Here, we asked how people acquire and accept assistance during adaptation to walking on a split-belt treadmill. Acquiring work from the treadmill and then accepting this work requires coordination between limbs and across different parts of the gait cycle, all while maintaining a steady pace to avoid drifting back or walking off the treadmill. We found that during adaptation, participants adopt asymmetric step lengths to acquire positive work from the treadmill and reduce the positive work by the legs. This reduction was associated with continuous reductions in metabolic cost over the course of 45 minutes of walking. Our findings support the idea that energetic cost, which can be reduced when learning to use external assistance, plays a critical role in shaping locomotor adaptation.

Previous studies have considered adaptation of locomotor patterns during split-belt walking to be complete once participants achieve symmetric step lengths (Day et al., 2018; Leech and Roemmich, 2018; Long et al., 2016; Mawase et al., 2017, 2012; Reisman et al., 2005; Roemmich et al., 2016). This is largely because people transition from a gait with marked step length asymmetries to taking steps of nearly equal lengths after 10 – 15 minutes of adaptation. Compared to the rapid changes in asymmetry observed during the first minutes of adaptation, the rate of adaptation slows markedly as adaptation continues beyond 15 minutes. However, the small changes in spatial and temporal measures of gait occurring after 15 minutes of adaptation continue for up to 45 minutes and are associated with reductions in positive work by the legs and energetic cost. Based on these results, we conclude that minimizing step length asymmetry, which was traditionally thought of as the goal of adaptation is perhaps better viewed as a point along the slower path toward a less energetically costly gait.

While it may seem intuitive that external assistance leads to a reduction in metabolic cost, providing participants with a source of external positive work does not imply that they will know how to use it for assistance. For example, participant 12 in our sample did not change their step lengths during adaptation (Supplementary Fig. 1) and they maintained the same rate of positive work with the fast leg through the entire 45 minutes of adaptation (Supplementary Fig. 2). Consequently, they did not reduce metabolic cost during adaptation (Fig. 6B). In participants who do learn how to take advantage of assistance, they learn to acquire positive work from the treadmill by first increasing forward limb placement of the fast leg, which increases negative work by the fast leg and this occurs within hundreds of strides. Then, they accept the positive work by the treadmill by reducing the positive work by the leg on the fast belt, but this process requires thousands of strides and had not ended even after 45 minutes. The time during the gait cycle when participants generate positive work is also crucial for reducing energetic cost (Donelan et al., 2002). Participants adjust the time when they generate positive work from single limb support during early adaptation to the step-to-step transition during late adaptation (Selgrade et al., 2017a). Generating positive work during the step-to-step transition at the same time as negative work is generated is less energetically costly as it decreases the cost of redirecting the body velocity (Donelan et al., 2002; Kuo, 2001). This learning problem of coordinating spatiotemporal, kinematic and kinetic processes to take advantage of external assistance appears to be complex enough complex enough that it requires the neuromotor system even longer than 45 minutes to solve.

We observed continuous changes in fast leg placement in front of the body, fast leg step length, and fast leg positive work throughout the 45 minutes of adaptation, whereas limb placement and work generated by the slow leg quickly reached a steady state. Specifically, participants increase the forward placement of the fast leg to acquire positive work from the treadmill and more gradually reduce positive work by the fast leg. Participants never increased positive work with either the fast or slow leg, indicating that they preferentially make gait modifications that reduce energetic cost and avoid strategies that might increase the amount of positive work performed by the legs. That the fast leg drives most gait adjustments is consistent with previous research showing that the leg on the fast belt accounted for the largest reductions in mechanical work (Selgrade et al., 2017a) and muscle activity (Finley et al., 2013) during adaptation.

A remaining question is, how long is required for full adaptation? Based on the two exponential model presented here, adaptation of step length asymmetry is near the predicted plateau at 45 minutes of split-belt walking. However, as evidenced by the fast leg positive work rate (Fig. 4C), adaptation of positive work rate has yet to achieve a stable plateau even after 45 minutes. Thus, adaptation of step length asymmetry does not mean that the entire adaptation process is complete even after ~2300 strides. While we cannot speculate on the amount of time required for complete adaptation, we know that it is markedly longer than is traditionally allotted. Similar to what has been observed with other types of adaptation studies such as reaching (Maeda et al., 2018), the time required for the slow processes involved in adaptation to plateau seems to be longer than what is traditionally allotted experimentally. This does not imply that adaptation cannot occur rapidly with other experimental manipulations. There is evidence that people adopt longer steps with the leg on the fast belt when walking on an inclined split-belt treadmill for 10 minutes (Sombric et al., 2019) or when adapting to belt speeds close to running for five minutes (Yokoyama et al., 2018). Given that these studies did not calculate mechanical work, and we do not know what the energetically optimal solution for the above studies would be, we cannot speculate if the rate of adaptation toward the energy optimal behavior occurred more rapidly than in the current study.

We found that the amount of experience did not affect the amount of retention of step length asymmetry, as indicated by the similar step length asymmetries measured during early post-adaptation following 15 and 45 minutes of adaptation. That the magnitude of step length asymmetry at early post-adaptation does not depend on prior exposure duration might be explained by task constraints: an initial asymmetry of ~0.30 with belts tied might be the largest asymmetry possible to avoid falling, walking off or drifting on the treadmill, but this remains to be determined. However, prolonged experience was associated with longer time constants for washout of the adapted pattern. Previous studies in reaching (Huang and Shadmehr, 2009) and walking (Roemmich and Bastian, 2015) have observed similar results for both the magnitude of the aftereffects and the duration of washout. After reaching in a force field, errors during error-clamp trials were similar after short versus long adaptation, yet errors persisted longer after adapting for more trials (Huang and Shadmehr, 2009). These findings combined with our own might be explained by a two state model of motor learning (Smith et al., 2006), where the time course of the decay of a motor memory depends on the number of adaptation trials, which here would correspond to the number of strides during adaptation. Thus, prolonged washout may reflect the state of the slower components of the adaptation process that are stabilized with practice.

## Conclusion

Prolonged adaptation to walking on a split-belt treadmill leads to continuous and gradual spatiotemporal, kinematic, and kinetic changes over different timescales, which result in a reduction in positive work by the legs and reduced energetic cost compared to what is traditionally observed after 15 minutes of adaptation. These findings demonstrate that participants learn to take advantage of the assistance provided by the treadmill, but this process requires much longer than is traditionally allotted during studies of adaptation. Learning to take advantage of external assistance is a complex problem since participants must first determine how to acquire assistance, and then learn how to take advantage of this assistance to reduce energetic cost. Thus, providing participants with enough time is crucial for the neuromotor system to adjust coordination and adapt the many different processes required to be able to exploit external assistance and reduce energetic cost. Together, these observations support the theory that locomotor adaptation is driven by energy optimization.

## Author Contributions

**Natalia Sánchez:** Conceptualization, Methodology, Software, Formal analysis, Investigation, Data curation, Writing—original draft, Writing—review & editing, Visualization, Project administration, Funding acquisition.

**Surabhi N. Simha:** Software, Validation, Writing—review & editing.

**J. Maxwell Donelan:** Methodology, Writing—review & editing, Supervision, Funding acquisition.

**James M. Finley:** Conceptualization, Methodology, Resources, Writing—review & editing, Supervision, Funding acquisition.

---

**Supplementary Figure 1.**
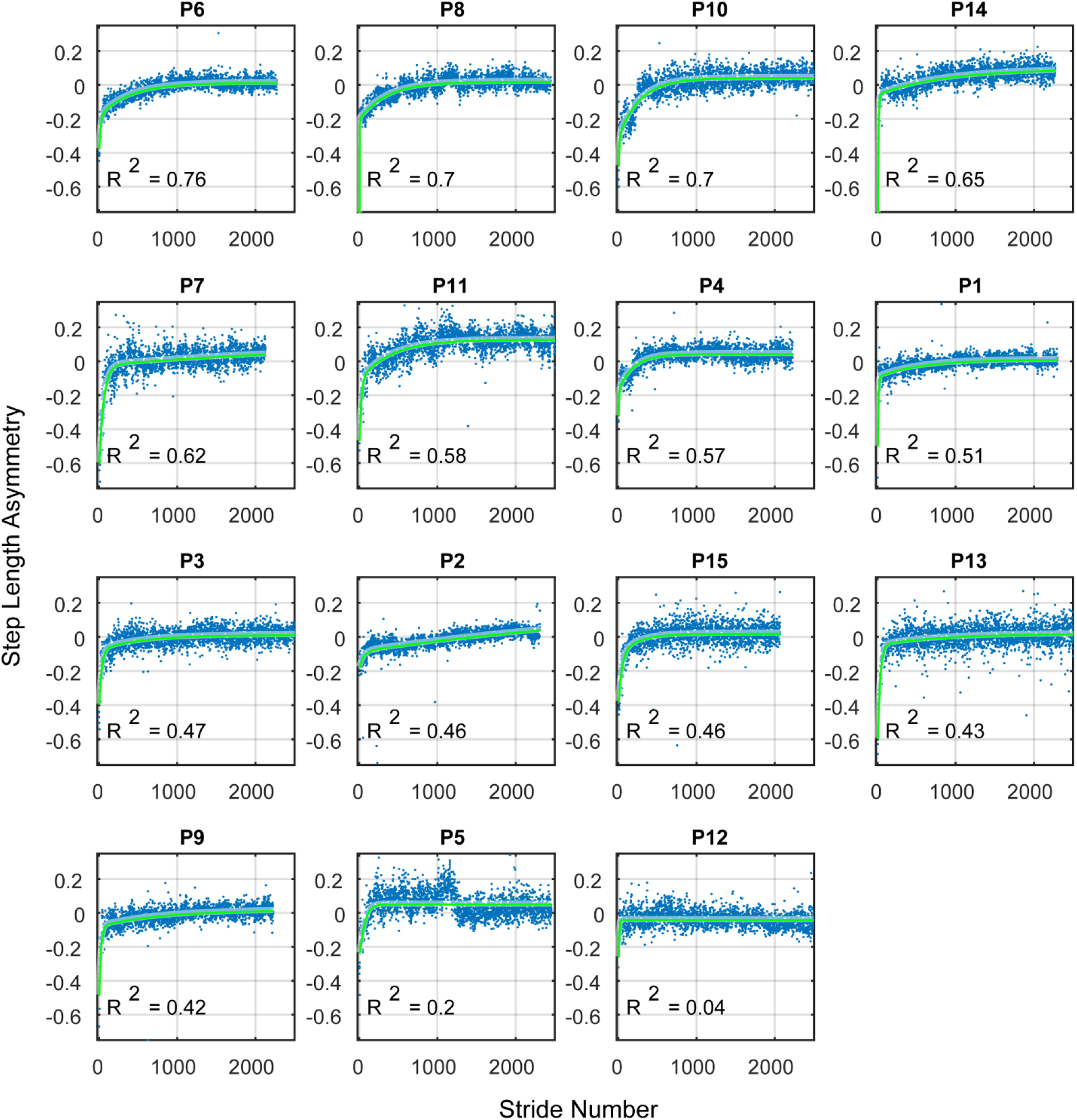
Individual fits using a two-exponential model for step length asymmetry sorted by R^2^. Step length asymmetry data was best fit using a two-exponential model. Fits are shown in green and raw data in blue.

**Supplementary Figure 2.**
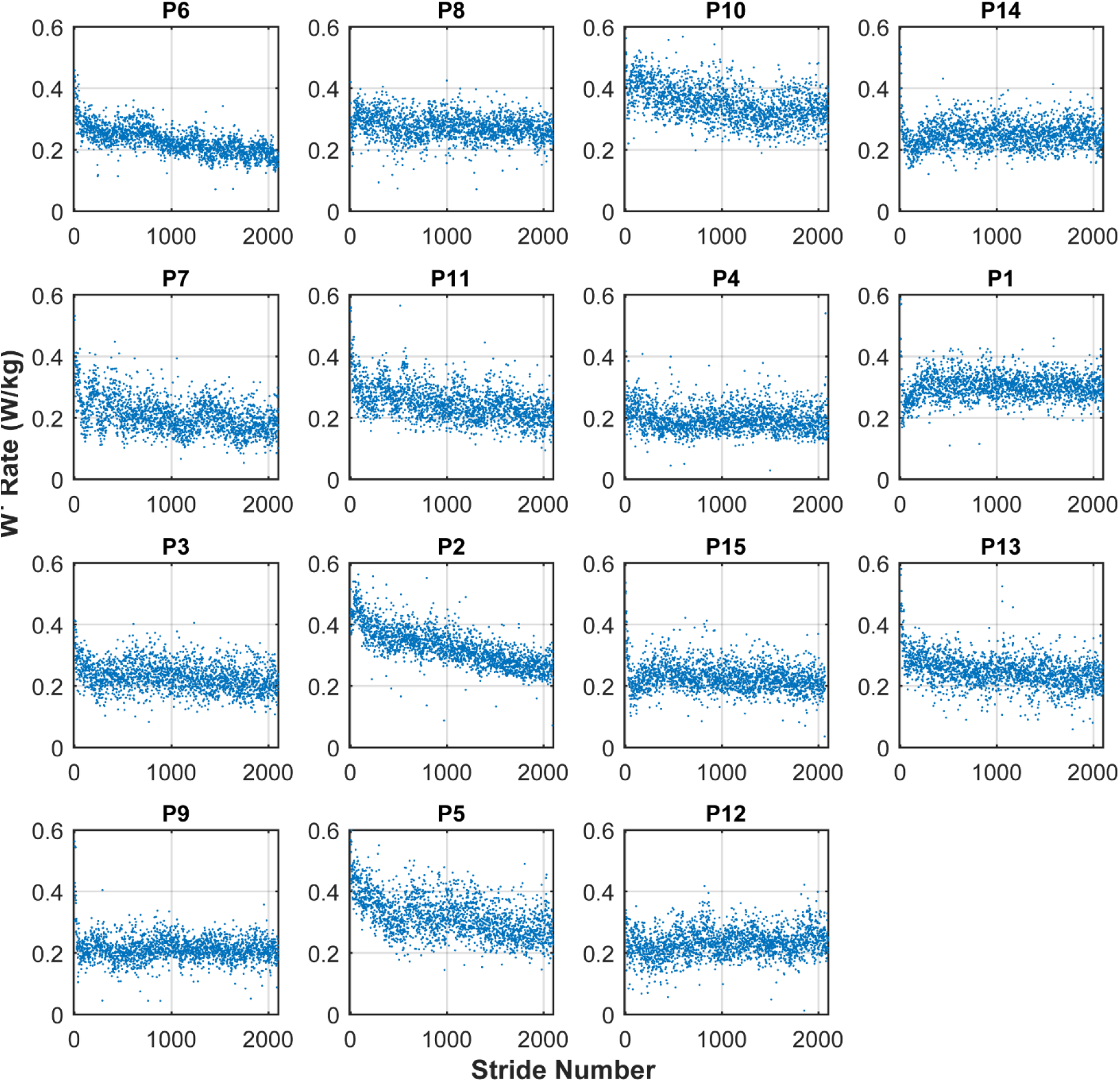
Individual data for positive work rate by the fast leg during adaptation. Participants are sorted as in Supplementary Fig. 1.

